# Spatial transcriptomics of compartmentalised inflammation in multiple sclerosis

**DOI:** 10.1101/2025.09.10.675210

**Authors:** Rachael Kee, Ger Mullan, Guillermo Lopez Campos, Gavin McDonnell, Valeriia Antsiferova, Michael John Dolan, Sam Loveless, Neil P. Robertson, Dessi Malinova, Owain Howell, Denise C. Fitzgerald, Michelle Naughton

## Abstract

Compartmentalised inflammation is a poorly understood aspect of multiple sclerosis (MS) that is associated with worse outcomes and represents an important therapeutic target. To gain deeper insight into compartmentalised inflammation, we have taken the approach of digital spatial profiling of the whole human transcriptome in areas of perivascular and meningeal inflammation and tertiary lymphoid-like structures (TLS) in MS central nervous system tissue. Critically, we had access to rare archival tissue obtained before the era of disease-modifying therapies, representing a natural history of disease. This analysis has identified differentially expressed genes in TLS compared to meningeal or perivascular inflammation. Pathway analysis highlighted that TLS signalling is dominated by B cell activity including active antibody secretion. Our data demonstrated the diversity of immunoglobulins and the prominence of IgG3- and IgG4-secreting cells in TLS. Intriguingly, pathway analysis suggests TLS may be hubs for viral (re)activation which warrants further investigation. These findings provide insight into the function of TLS in MS disease pathogenesis and reveal unique immune signatures that may support biomarker development to predict which patients harbour TLS in life.

## Introduction

Multiple sclerosis (MS) is a chronic, immune-mediated, demyelinating disease of the central nervous system (CNS). Whilst disease-modifying treatments (DMTs) have improved patient outcomes through effective suppression of inflammatory lesions, they have limited effectiveness against disease progression. Progression independent of relapse activity (PIRA) and/or new disease activity detected by magnetic resonance imaging, underlies the majority of MS disability.^1–3^ Compartmentalised inflammation is hypothesised as a possible pathological driver of PIRA.^4^ It refers to inflammation in meningeal and perivascular spaces and is associated with increased innate immune activation in both lesioned and non-lesioned areas. Imaging and neuropathological studies detected an outside-in gradient of pathology on all brain surfaces exposed to cerebrospinal fluid (CSF).^5–7^ Meningeal inflammation is associated with a surface-in gradient of microglial activation and cortical loss of neurons, astrocytes and oligodendrocytes.^8–12^ Tertiary lymphoid-like structures (TLS) are a feature of the most severe grade of compartmentalised inflammation.^13^ In comparison to TLS associated with tumours and autoimmune conditions, TLS in MS are more loosely organised aggregates, often lacking defined T and B cell zones and markers of germinal centre reactions.^4^ Nonetheless, they are associated with more severe MS, incurring earlier age of onset, disability and death.^10^ We report the first whole transcriptome characterisation of TLS in post-mortem MS cases of natural disease history, identifying differentially expressed genes in TLS compared to surrounding meningeal inflammation. Signalling pathway analysis pointed to active antibody-secreting B cells in TLS. This provides important insight to the role of TLS in MS disease pathogenesis and may reveal unique immune signatures to better predict which patients harbour TLS in life.

## Materials and methods

Detailed materials and methods are provided in the Supplementary Methods.

### Screening for presence of TLS

Meningeal and perivascular inflammation in 53 cases in the Dame Ingrid Allen (DIA) tissue collection (Supplementary Table 1) was graded using previously published criteria.^10,14^ Cases harbouring substantial (^+++^) cellular aggregates were screened for the presence of CD3^+^ T cells and CD20 B^+^ cells. Four TLS^+^ cases were identified from the DIA tissue collection and five TLS^+^ cases from the MS Society Tissue Bank.

### Spatial transcriptomics

Nanostring GeoMx Digital Spatial Profiler whole transcriptome analysis was undertaken through their Technology Access Program (Nanostring Technologies, Inc, Seattle, USA) on FFPE tissue from two TLS^+^ post-mortem cases from the DIA tissue collection. Regions of cortex and compartmentalised inflammation containing clusters of CD3^+^ T cells and CD20^+^ B cells in TLS, meninges and perivascular spaces of white (WM) and grey matter (GM) were defined as regions of interest for analysis (Supplementary Figure 1). RNAscope HiPlex V2 assay was used as per manufacturer’s instructions to detect *IGHG, IGHM, IGHA* and RNAscope negative and positive control probes in one TLS^+^ post-mortem case (Supplementary Table 2). Images were acquired on a Stellaris 5 confocal microscope and processed in Fiji ImageJ.

## Results

### Molecular profiles of CNS compartmentalised inflammation

174 FFPE cortical tissue blocks from 53 post-mortem cases from a natural history cohort were graded for compartmentalised inflammation and screened for presence of TLS (Supplementary Tables 1-2). In agreement with previous findings^10^, cases with substantial meningeal inflammation associated with shorter disease duration and younger age at death (Figure 1A-E). Four TLS^+^ cases were identified.

**Figure 1.**
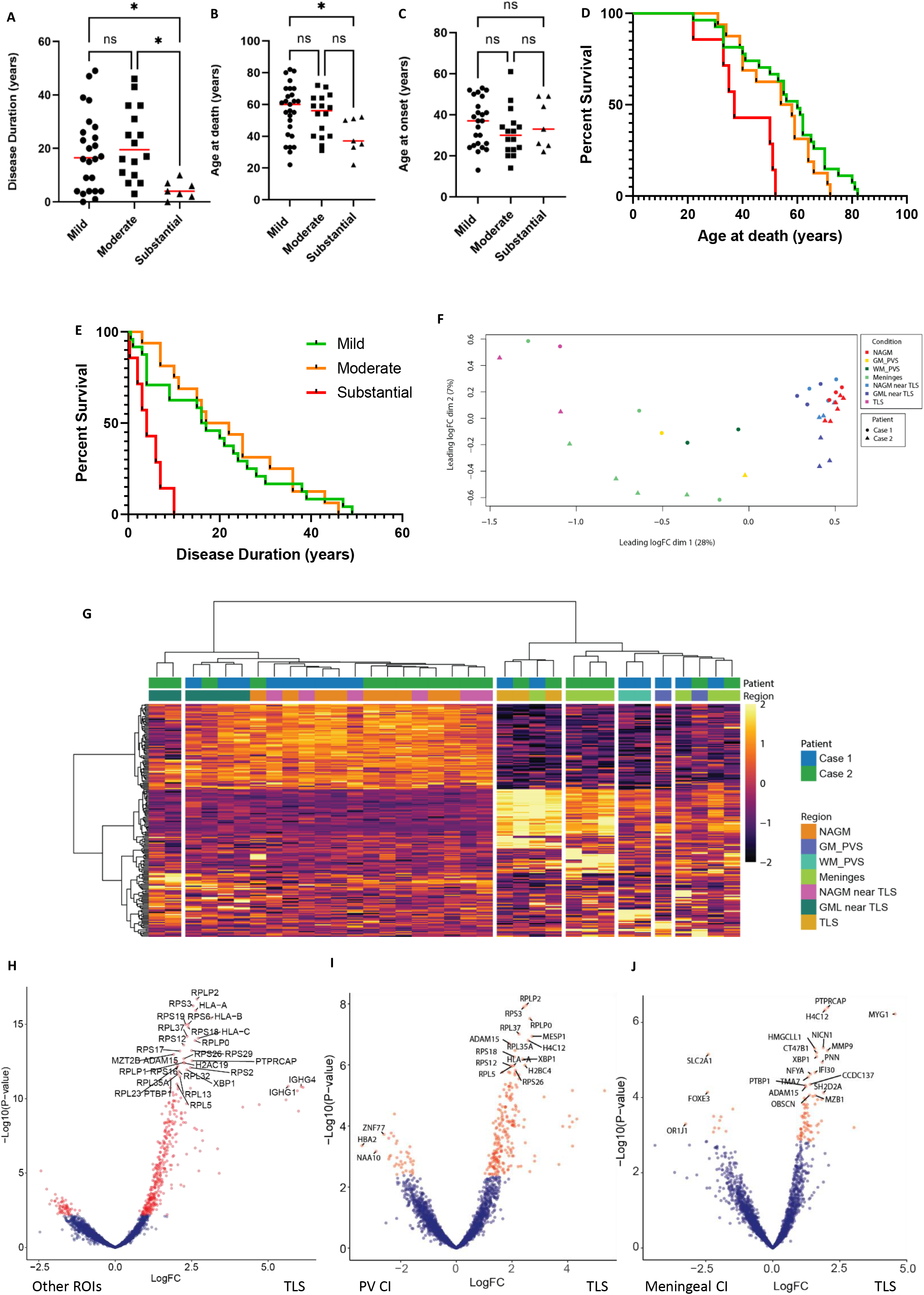
Transcriptomics of compartmentalised inflammation in MS. Substantial levels of meningeal inflammation in MS were associated with a more severe disease course. Age at onset (A), death (B, D) and disease duration (C, E) was compared between cases with mild, moderate or substantial levels of meningeal inflammation. Cases that had substantial meningeal inflammation were associated with a shorter disease duration (D) and younger age at death (E). MDS plot of regions of interest (F). Unsupervised hierarchical clustering of log2 transformed values were used to compare expression levels between regions (G). Volcano plot showing the differentially expressed genes between TLS and all other ROIs (H), TLS vs regions of meningeal inflammation (I) or perivascular inflammation (G). Abbreviations: NAGM, normal-appearing grey matter; GML, grey matter lesion; PVS: perivascular space; TLS: Tertiary lymphoid-like structure; WM, white matter.

We performed whole transcriptome analysis on areas of compartmentalised inflammation (perivascular inflammation, meningeal inflammation, TLS) and cortex (normal-appearing grey matter and demyelinated grey matter lesion) of two TLS+ cases (post-mortem interval of Case 1: < 12h, Case 2: 24h). Hierarchical clustering demonstrated separate clusters for cortical regions and for TLS and most other regions of compartmentalised inflammation (Figure 1F,G). Many of the most abundantly expressed genes related to immunoglobulins (*IGHG1-4, IGHM, IGKC, IGLL5, IGHA1, JCHAIN)* indicating wide B cell diversity within TLS (Fig 1H, Supplementary Table 6). Genes encoding IgG3 and IgG4 were the highest expressed subclasses. Other abundantly expressed genes in TLS included genes associated with ribosomes and protein synthesis, class I and II antigen presentation (*CD74, B2M, HLA-A, HLA-B, HLA-C, HLA-DRB1, HLA-E*), complement (*C1QA, C1QB*), genes involved in assembly and secretion of immunoglobulins (*MZB1* and *XBP1*), chemokine *CXCL9* and cytokines *CCL5* and *CCL19* associated with B and T cell trafficking, as well as *IFITM1*, an IFN-induced antiviral protein.

Gene ontology of biological and molecular functions of TLS demonstrated strong enrichment for genes associated with transcription, protein synthesis/translation and antigen processing (Supplementary Figure 2A,B). In addition, Reactome analysis detected “viral mRNA translation”, “nonsense-mediated decay” and “response of EIF2AK4 to amino acid deficiency” (cellular response to stress/starvation/viral infection), whilst KEGG identified strong enrichment for pathways relating to viral infections and autoimmunity (Supplementary Figure 2C,D).

### Differential gene expression between TLS and other CNS inflammatory regions

Differentially expressed genes between TLS and perivascular inflammatory compartments again included genes associated with immunoglobulins and their secretion, class I and class II antigen presentation, and chemokines and cytokines associated with B and T cell trafficking, suggesting that TLS are regions of higher organisation of B and T cells in comparison to CNS perivascular inflammatory regions (Figure 1I, Supplementary Table 7).

The most differentially expressed genes between TLS and other inflammatory meningeal regions included *MYG1*, an RNA exonuclease that mediates nucleo-mitochondrial crosstalk, *PTPRCAP*, a key regulator of T and B lymphocyte activation, and *MMP9* which is upregulated in CSF of MS patients.^15,16^ Other significantly increased genes again included *MZB1* and *XBP1* required for plasmablast differentiation and suggestive of germinal centre activity,^17,18^ as well as *IFI30* involved in class II antigen presentation, *CCL5* cytokine also known as RANTES, and *SRPRB* which is highly expressed in plasma cells (Figure 1J). Intriguingly, *ADXDN1* and *CT47B1* were also revealed as differentially expressed genes though these have not previously been associated with immune or CNS cells. Overall, this gene expression profile suggests that, in comparison to other meningeal regions harbouring diffuse collections of immune cells, TLS may be sites of organised B cells and a source of immunoglobulin-secreting plasma cells. In contrast, few significant differences were observed between perivascular and meningeal regions of compartmentalised inflammation (Supplementary Figure 2E, Supplementary Table 8)

### IgG3 and IgG4 profiles in regions of compartmentalised inflammation

The highest expressed genes detected in TLS, namely immunoglobulins, were validated with HiPlex RNAscope in one TLS case (Supplementary Figure 3). While the highest gene expression was recorded for IgG subtypes in TLS, a higher proportion of TLS cells were positive for *IGHM* (25%) versus *IGHG* (17%). The percentages of cells positive for *IGHG* and *IGHA* mRNA were similar though levels of gene expression for *IGHG* were much higher. All three immunoglobulin subtypes showed similar patterns of distribution across compartments; TLS and perivascular spaces in WM or adjacent cortex contained higher levels of each subtype, whereas other areas of meningeal inflammation showed few positive cells, and none were detected in cortical perivascular spaces distal to TLS (Supplementary Figure 3A-C).

Next, we characterised IgG and the highest expressed subsets, IgG3 and IgG4, at protein level in TLS versus other regions of compartmentalised inflammation in a wider TLS^+^ MS cohort (n=5-8 per region). Whilst cells were most densely concentrated in TLS, the percentages of IgG^+^, IgG3^+^ cells or IgG4^+^ cells were not significantly different between regions (Figure 2A-D, Supplementary Figure 4). In TLS, significantly higher percentages of IgG4^+^ than IgG3^+^ cells were observed (mean 9.75% vs 2.84%) (Figure 2E-G), a trend observed in all inflammatory regions (Figure 2C-D). When TLS^+^ cases were stratified by short or long disease duration (<4 years, n=3. >10 years, n=2), this trend amplified over time (short disease: mean 3.5% IgG3^+^ vs 5.6% IgG4^+^. Long disease: mean 1.8% IgG3^+^ vs 15.9% IgG4^+^) (Figure 2G).

**Figure 2.**
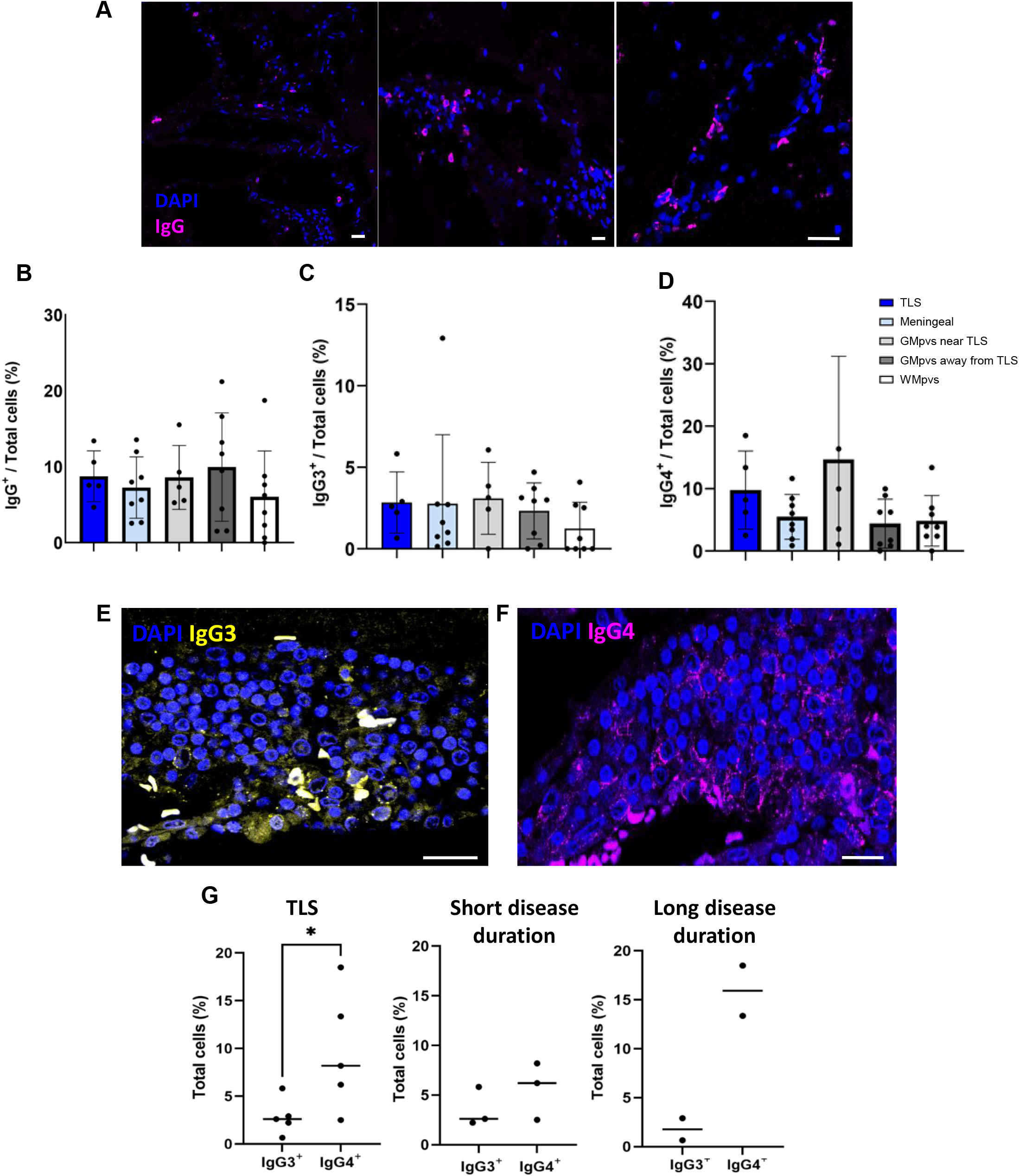
IgG, IgG3 and IgG4 detection in post-mortem MS cohort. The proportion of mean IgG^+^ cell counts were not statistically different across regions of interest (A). Variation in IgG^+^cell density was observed across post-mortem cases. Detection of IgG^+^ cells in inflammatory meningeal regions and GM perivascular spaces (B). The proportion of TLS cells positive for IgG3 (C) and IgG4 (D) was similar across all regions of interest and no statistically significant difference was observed. Representative images showing IgG3^+^ cell detection (E) and IgG4^+^ cell detection (F) in TLS of the same case. IgG4^+^ cell densities were higher in TLS compared with IgG3^+^ cell densities (G) n=5, *p=0.0340 unpaired two-tailed Student’s t-test after arcsin transformation. In cases with short disease duration (<4 years) IgG3 and IgG4 cell densities in TLS were similar. In longer disease duration (>10 years) cases there was a trend of a higher IgG4^+^ cell density in TLS compared with IgG3^+^ cell density (G). Scale bar = 20 µm. TLS and GMpv near TLS n=5, non-TLS meningeal, GMpvs away from TLS and WMpvs n=8. Abbreviations: TLS; tertiary lymphoid-like structure, GMpvs; grey matter perivascular space, WMpvs; white matter perivascular space.

### IgG3^+^ signal co-localised with CD38^+^, CD138^+^ or CD38^+^CD138^+^ cells in TLS

Plasma cells are characterised by immunoglobulin secretion and may be CD38^+^CD138^high/low^, while CD38^+^/CD138^-^ can be representative of germinal centre B cells.^19,20^ In TLS, approximately 60% of IgG3^+^ cells co-localised with CD38, including 20% CD38^+^CD138^+^ plasma cells (Figure 3). Very few (∼2%) co-localised with CD138 single positive cells. In contrast, the majority of IgG4^+^ cells did not co-localise with CD38^+^, with very few double-positive CD38^+^CD138^+^ cells or CD138^+^ cells. In TLS, approximately 30% of cells were CD38^+^IgG4^+^, less than 5% of cells were CD38^+^CD138^+^IgG4^+^ and none were CD138^+^IgG4^+^ (Figure 4A-E).

**Figure 3:**
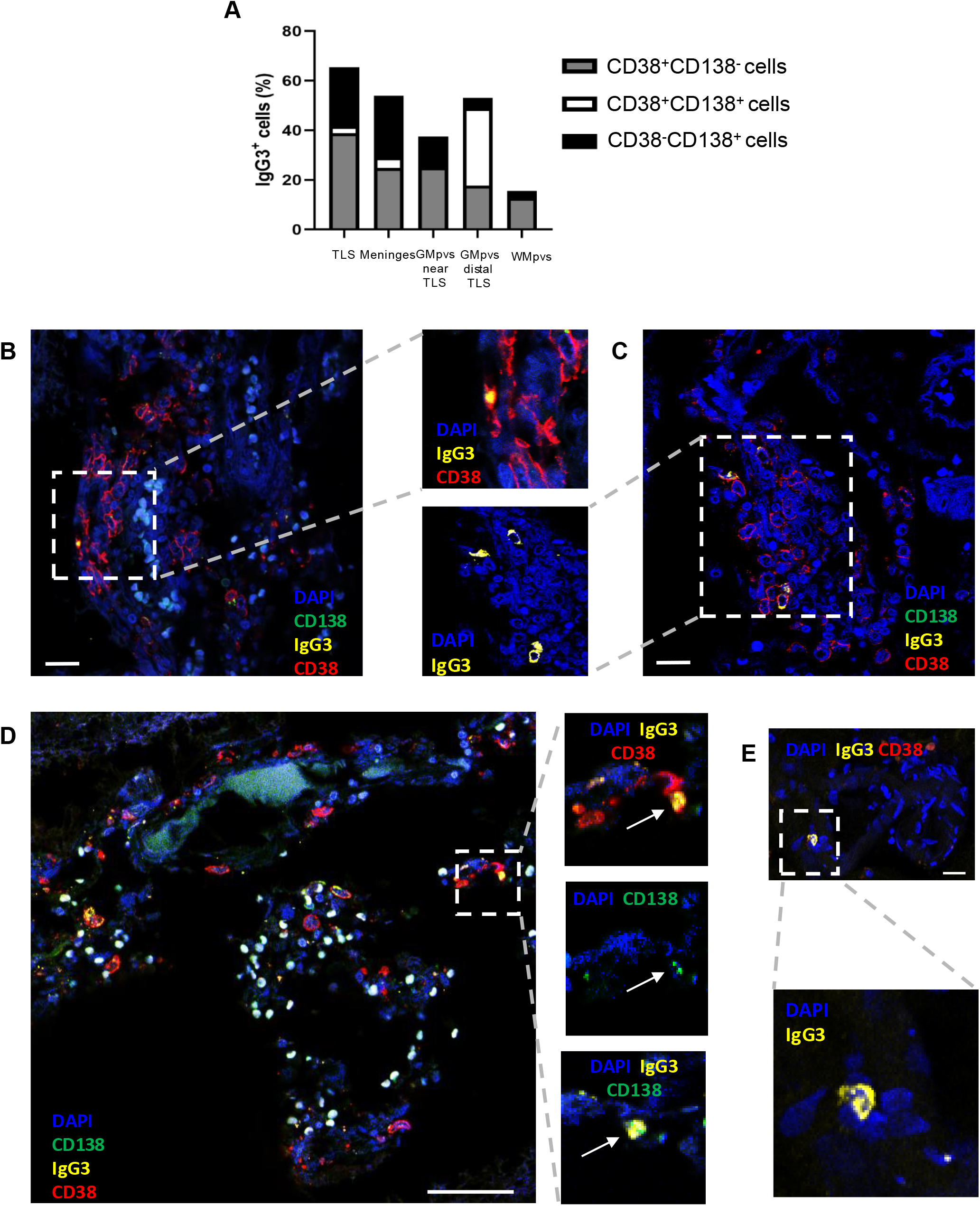
IgG3 co-localisation with CD38^+^, CD138^+^ and CD38^+^ CD138^+^cells. The majority of IgG3^+^ cells co-localised with CD38^+^, CD138^+^or CD38^+^CD138^**+**^cells in TLS, non-TLS meningeal regions and GM perivascular spaces (A). Representative images from two TLS cases showing co-localisation of IgG3^+^cells with CD38^+^cells (B, C). CD38^+^IgG3^+^cells and CD38^+^ CD138^+^ IgG3^+^ cells (arrows) were also observed in meningeal spaces (D). In WM perivascular spaces IgG3^+^ cells were often not co-localised to CD38 or CD138 (E). Scale bar = 20 µm (B, C, E), 50 µm (D). Abbreviations: TLS; tertiary lymphoid-like structure, GMpv; grey matter perivascular, WMpv; white matter perivascular. TLS and GMpv near TLS n=5, non TLS meningeal, GMpv distal from TLS and WMpv n=8.

**Figure 4:**
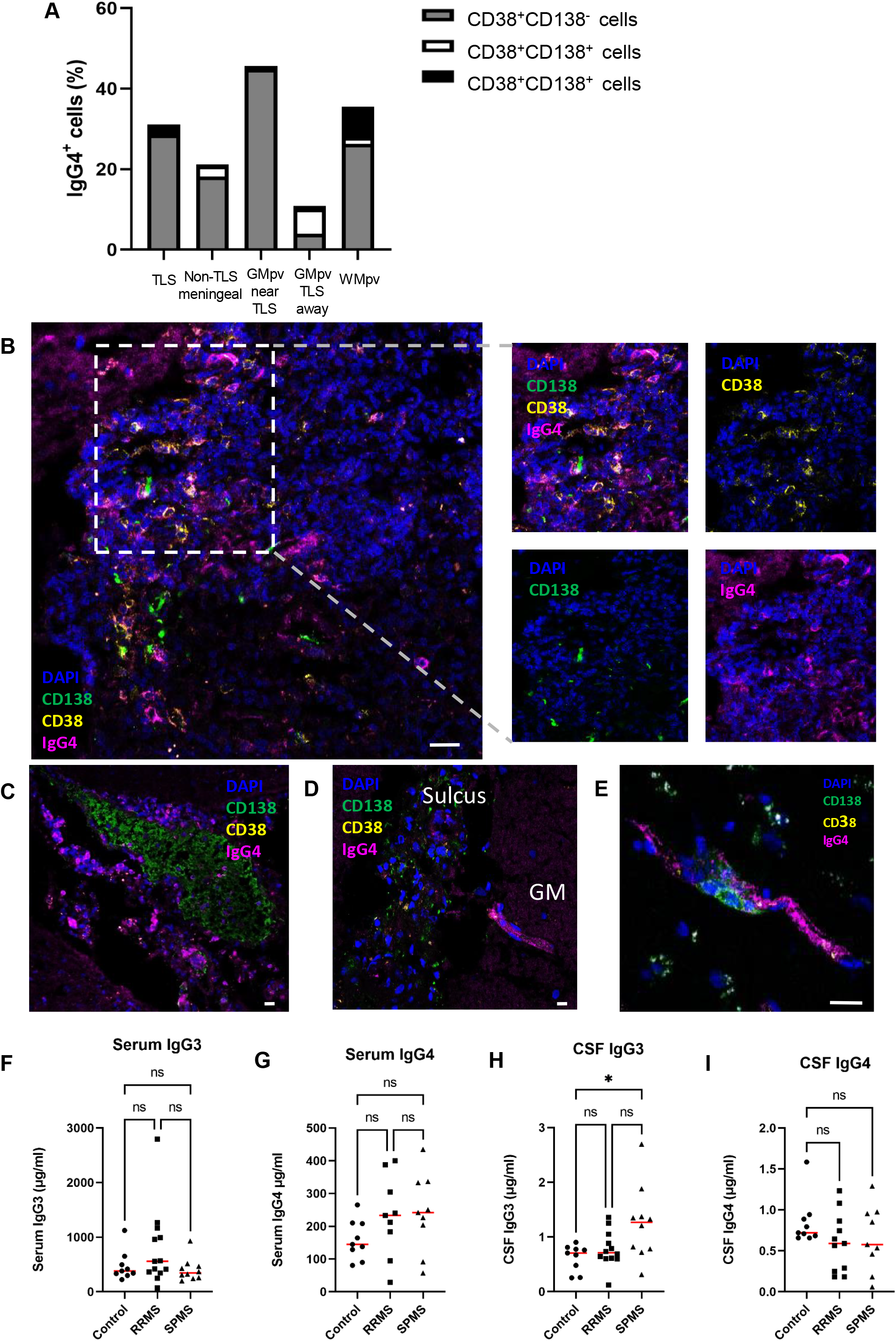
IgG4 co-localisation with CD38^+^, CD138^+^ and CD38^+^ CD138^+^ cells. The proportions of CD38^+^ IgG4^+^, CD138^+^ IgG4^+^ and CD38^+^ CD138^+^ IgG4^+^ cells across the ROIs was variable (A). Very few CD138^+^ IgG4^+^ and CD38^+^ CD138^+^ IgG4^+^ cells were observed. Most IgG4^+^ cells did not co-localise with CD38^+^, CD138^+^, or CD38^+^ CD138^+^ cells. The detection of CD38^+^IgG4^+^cells was observed in the periphery of a TLS from case MS601 (B) and interspersed throughout the TLS in case MS584 (C). The appearance of cells positive for IgG4 and secreted IgG4 was observed in GM perivascular spaces adjacent to the meninges and near to TLS (D). WM perivascular spaces also had the appearance of secreted IgG4, sometimes in close proximity to CD138^+^cells (E). Scale bar = 20 µm. Serum and CSF IgG3 and IgG4 levels in cohort that remained RRMS at follow up (RRMS cohort, n=13) and those that developed SPMS (SPMS cohort, n=10) and non-MS controls (n=9) (F-I). Abbreviations: TLS; tertiary lymphoid-like structure, GMpv; grey matter perivascular, WMpv; white matter perivascular. TLS and GMpv near TLS n=5, non TLS meningeal, GMpv away from TLS and WMpv n=8.

### Initial CSF IgG3 levels significantly raised in those that develop SPMS at follow-up

Matched serum and CSF samples were used to investigate whether initial levels of IgG3 and IgG4 differed between relapsing-remitting (RRMS) patients who remained RRMS at follow-up or converted to secondary progressive MS (SPMS), and other neurological controls. At lumbar puncture, the RRMS cohort had median Expanded Disability Status Scale (EDSS) 2 and the SPMS cohort had 3.25. At follow-up, all of the RRMS cohort remained <4.0, whereas EDSS ranged from 5-8.5 in the converted SPMS cohort (Supplementary table 5).

Serum IgG3 and IgG4 levels were comparable across all cohorts (Figure 4F-I). CSF IgG3 levels were significantly higher in those that later converted to SPMS compared to the control cohort, in contrast to those who remained RRMS at follow-up. CSF IgG4 levels were comparable between cohorts. There was no significant correlation between disease duration and serum or CSF IgG3 or IgG4 (Supplementary Figure 5).

## Discussion

Compartmentalised inflammation is a poorly understood aspect of MS associated with worse outcomes, a finding which we validated in this study cohort. Critically, we had access to rare archival tissue collected before the advent of disease-modifying therapies, representing natural history of disease. Although technically challenging, we demonstrate the feasibility of digital spatial profiling with tissues dating from the 1960s.

We uncovered TLS gene signatures featuring upregulated immunoglobulins and chemokine production, and specifically upregulated genes relative to other sites of meningeal inflammation including *MYG1, PTPRCAP* and *MMP9*. This presents the possibility of using these targets as indicators of TLS and warrants analysis in CSF in future studies. We conducted pathway analysis to gain a more holistic understanding of potential TLS biological functions and activity. This was dominated by pathways associated with translation, aligning with the exceptional demand on protein synthesis machinery of antibody-secreting B cells, and with immunoglobulins being the most robustly expressed genes. *XBP1* and *MZB1* expression, abundant immunoglobulin transcripts, and CD38/CD138 co-localisation are consistent with plasma cell differentiation and antibody secretion. These data support the prominence of B cells in TLS reported earlier, and we confirm that expression profiles and signalling pathways of TLS are dominated by B cell function and development. Given potent therapeutic effects of B cell-depleting monoclonal antibodies on peripherally-driven inflammation in MS, it will be important to determine whether these interventions impact the formation, stability or function of TLS and other features of CNS compartmentalised inflammation and associated aggressive disease courses in MS. Given the very low CNS penetrance of monoclonal antibodies, our study suggests that a B cell-targeting strategy active within the CNS may be optimal.

Intriguingly, pathway analysis identified virus-associated signalling as some of the most enriched pathways in TLS. As this analysis is restricted to the human transcriptome, this is largely due to differences associated with hijacking of host translational machinery. *PABPC1* and a multitude of ribosomal targets were upregulated in TLS in our dataset, including *RPL22* which can bind EBV-specific proteins.^21^ While this could potentially represent viral (re)activation in infected immune cells, these genes can overlap with protein synthesis pathways employed by B cells for antibody secretion,^22^ while some such as *RPL22* have a crucial role in B cell development.^23^

We validated key targets at single-cell level using HiPlex RNAscope. This indicated substantially higher IgG mRNA expression per cell than IgM or IgA. At the protein level, IgG3^+^ and IgG4^+^ cells associated with germinal B cell and plasma cell markers CD38 and CD138. In TLS, there were significantly higher numbers of IgG4^+^ cells in comparison to IgG3^+^ cells, with a trend towards greater discrepancy in long-duration cases. While this should be validated in larger, covariate-adjusted cohorts, we also observed significantly higher CSF IgG3 levels in patients that later converted to SPMS compared to controls, in contrast to those who remained RRMS at follow-up. IgG3 is a potent, pro-inflammatory antibody with a short half-life associated with viral infections and autoimmune disorders. It has the highest affinity for the complement protein C1q^24^ and is a potent activator of the classical complement pathway, which was also detected in our transcriptomic dataset and is associated with compartmentalised inflammation and cortical pathology in MS.^25^ The IgG4 subclass is associated with chronic antigen exposure in non-infectious settings, e.g. allergy. Both these subclasses are much less abundant in blood compared to IgG1 and IgG2. Nonetheless, Trend *et al*. found a correlation between serum IgG3 and time to MS diagnosis in clinically-isolated syndrome (CIS) patients; IgG3 levels increased as time to MS diagnosis drew nearer.^26^ This group showed the same correlation with higher IgG3^+^ B cells detected in CIS patients closer to relapse.^27^ These IgG3^+^ B cells expressed CXCR5, the receptor for CXCL13. It would be interesting to determine whether IgG3^+^ B cells also express CXCR5 to home to TLS in the CNS, and are associated with complement-mediated cortical pathology in MS.^25^ This provides further basis to investigate IgG3^+^ cells as potential therapeutic targets and biomarkers of disease activity.^27^

A limitation of the study was sample size of TLS^+^ cases in our archive. Only two cases underwent spatial transcriptomic profiling; disease duration was 6 years for case 1 and 4 months for case 2. While such small samples limit the representation of MS more generally, gleaning insight into early stages of MS in a natural disease cohort is a powerful addition to the field. The most highly expressed targets were validated using a larger cohort from the MS Society Tissue Bank. These findings should be investigated in a larger cohort, including in post-DMT era tissues, to determine if contemporary treatments affect TLS formation or functions.

In conclusion, to our knowledge this is the first report of TLS molecular profiles in MS across the whole human transcriptome. The dominance of B cell function in core TLS activity is highlighted by our findings of diverse immunoglobulins and prominence of IgG3 and IgG4-secreting cells in TLS. Pathway analysis points to viral-associated cellular responses within TLS immune cells, suggesting that TLS may be hubs for viral (re)activation which requires further investigation. The identification of secreted factors in TLS transcriptomic profiles points towards putative soluble biomarkers that could inform the stratification of cohorts with TLS and indeed responsiveness to future treatments targeting TLS formation and function. Collectively, our findings provide new insight into CNS compartmentalised inflammation that may support future biomarker development and therapeutic targeting of CNS-specific pathogenic mechanisms of MS.

## Acknowledgements

For tissue samples and associated clinical and neuropathological data supplied by the MS Society Tissue Bank (Imperial College London), funded by Multiple Sclerosis Society, registered charity 207495. Biosamples were obtained from the Welsh Neuroscience Research Tissue Bank which is partly funded by Health and Care Research Wales (Advanced Neuro Therapies Unit). Other investigators may have received samples from the same subjects.

