## Supplementary Material for "Spatial transcriptomics of compartmentalised inflammation in multiple sclerosis"

### Supplementary Materials and Methods

#### Ethics statement

Post-mortem tissues were sourced from the Dame Ingrid Allen (DIA) Tissue Collection (under ethical approval by the QUB Faculty Research Ethics Committee:14.57v4, MHLS 21_119) and the UK MS Tissue Bank (National Research Ethics Committee:18/WA/0238).

Samples of paired serum and CSF in the Welsh Neuroscience Research Tissue Bank was approved by the Wales REC3 National Research Ethics Service [19/WA/0058].

#### Spatial transcriptomics

Nanostring GeoMx Digital Spatial Profiler (DSP) whole transcriptome analysis was undertaken through their Technology Access Program (Nanostring Technologies, Inc, Seattle, USA). FFPE tissue sections from two TLS$^+$ post-mortem cases from the DIA tissue collection stained for MBP, CD3 and CD20. Regions of cortex and compartmentalised inflammation containing clusters of CD3$^+$ T cells and CD20$^+$ B cells within the meninges, perivascular spaces and TLS were defined as regions of interest (ROIs).

11 of 48 ROIs were removed due to low signal (largely affecting smaller ROIs). Sequencing metrics confirmed sufficient depth to detect low-expression targets and identify outliers. All 37 ROIs had high levels of sequencing saturation and detected a total of 18,676 targets. Differential expression analysis was performed using “edgeR” (v4.2)[1]. Genes were filtered using the filterByExpr function using default parameters, which we confirmed removed the negative control probe. We performed a TMM normalization, dispersion estimation and differential expression. We performed two parallel analyses: one comparing each region versus all others, and a second using individual contrasts. For the latter, we were interested in three contrasts: 1) TLS versus Meninges, 2) TLS versus Perivascular and 3) Meninges versus Perivascular. For these latter two contrasts, we pooled all perivascular samples, regardless of white or grey matter source. We used clusterprofiler (version 3.14.3)[2] enrich family of functions (“enrichGO”, “enrichPathway” and “enrichKegg”) for all over representation analysis.

#### Immunofluorescence

FFPE tissue sections were baked, dewaxed and rehydrated through a graded series of ethanol solutions prior to antigen retrieval. Fresh-frozen (FF) post-mortem brain sections were fixed in 10% neutral buffered formalin and did not go through antigen retrieval. Tissue sections were blocked and primary antibody solutions (supplementary table 4) incubated overnight at 4°C. Secondary antibodies were incubated for 30 minutes at room temperature in the dark. DAPI (1:10000) was used for nuclear stain. Slides were mounted with ProLong Gold. TrueBlack® Lipofuscin Autofluorescence Quencher (Biotium, 23007) was applied to FF tissue only, prior to the blocking step.

Images were acquired on a Stellaris 5 confocal microscope. The QuPath 'cell detection' tool was used to quantify nuclei counts. Immunofluorescent cells were counted manually and validated by a blinded, second counter. The proportion of positive cells per ROI were reported. Co-localisation of IgG3 and IgG4 with $CD38^+$ and/or $CD138^+$ cells was reported as a proportion of total $IgG3^+$ or $IgG4^+$ cells per ROI respectively. Representative images were processed in Fiji ImageJ.

#### ELISA

All cases with paired serum and CSF in the Welsh Neuroscience Research Tissue Bank were searched for a diagnosis of RRMS that had transitioned to SPMS at follow-up (n=10). Date of lumbar puncture was matched as close to initial symptom onset as possible. Cases with a diagnosis of RRMS who remained RRMS at last follow-up were matched to age, sex and disease duration of the SPMS group (n=13). The non-MS control group (n=9) were age-sex matched to the RRMS and SPMS cohort (supplementary table 5). ELISAs were performed according to manufacturer's instructions for serum and CSF IgG3 and IgG4 (Thermofisher, 991000) and analysed by 4-parameter curve fit.

#### Statistical Analysis

Data were checked for normal distribution using the Shapiro-Wilk test. Parametric datasets used unpaired Student's t-tests for comparisons of two groups, and one-way ANOVA with Tukey's multiple comparisons test for more than two groups. Statistical tests of non-parametric datasets used Mann-Whitney U-test for comparisons of two groups, and Kruskal-Wallis with Dunn's multiple comparison test for more than two groups. Percentage data underwent arcsin transformation prior to normality testing. To test correlation between two groups, the Pearson or Spearman's correlation test was used for parametric or non-parametric data respectively. GraphPad Prism (Version 9, GraphPad Inc.) was used for statistical analysis and graphing of data.

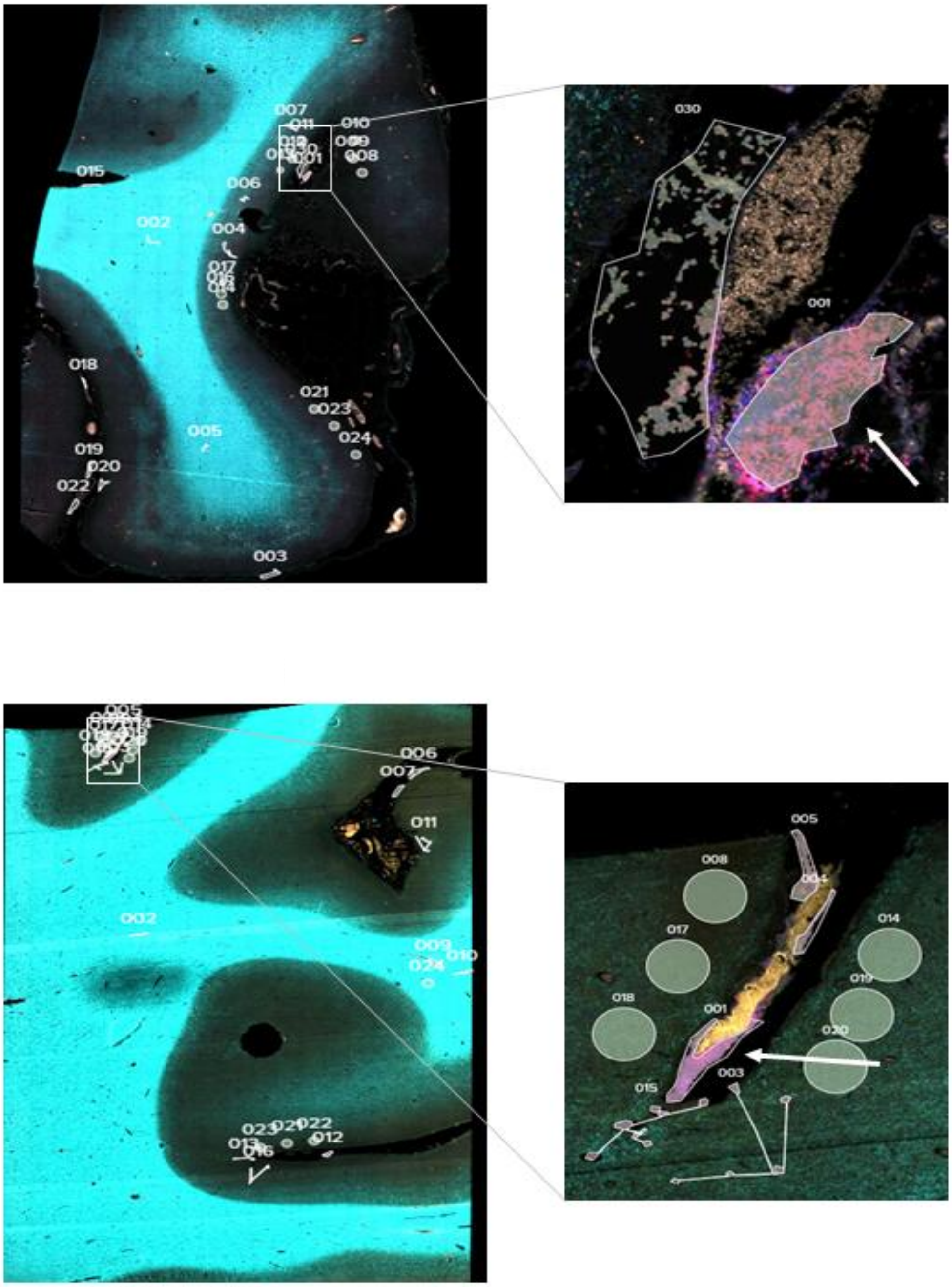

**Supplementary Figure 1: Nanostring digital spatial profiling of 2 post-mortem cases from the Dame Ingrid Allen collection.** Digitalised whole slide scans with regions of interest (ROI) that were selected for whole transcriptome analysis. Tertiary lymphoid-like structures (TLS, indicated by white arrows) and areas of meningeal and perivascular inflammation were detected by the presence of $CD3^{+}$ and $CD20^{+}$ cells and manually traced as ROIs. Other ROIs in the grey and white matter were selected with circles (not included in current analysis).

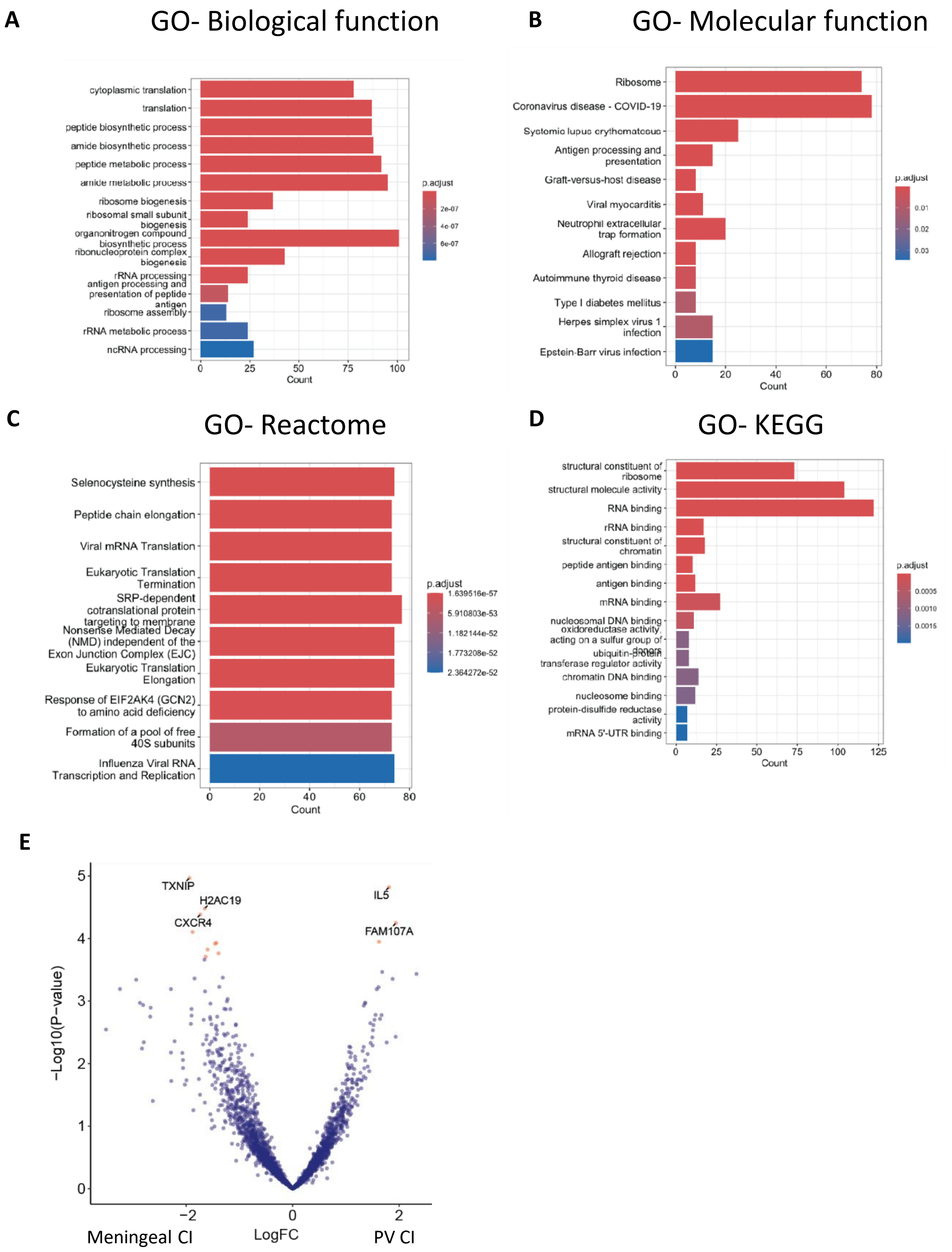


**Supplementary Figure 2. Pathway analysis of TLS gene expression.** Bar chart of top enriched terms from the Reactome_2022 gene set library. The top 10 enriched terms for the input gene set are displayed based on the -log10(p-value), with the actual p-value shown next to each term. The term at the top has the most significant overlap with the input query gene set.

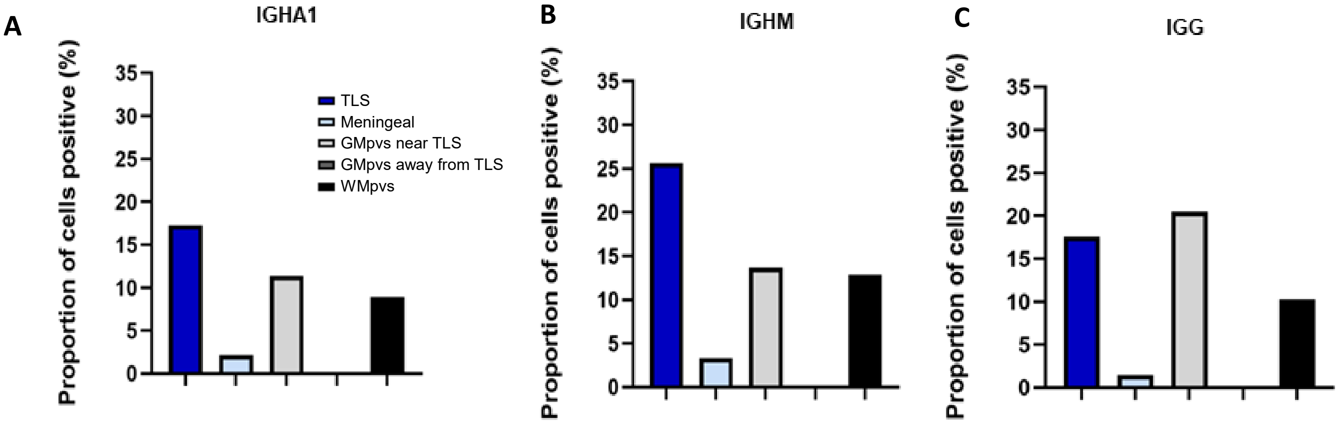


**Supplementary Figure 3. Profile of immunoglobulin subtypes in regions of compartmentalised inflammation.** Validation was undertaken of mRNA targets identified by transcriptomics in a single case using HiPlex RNAscope. Proportion of positive cells is reported for detection of mRNA transcripts of (A) *IGHA* (B) *IGHM* (C) *IGHG*. Abbreviations: TLS; tertiary lymphoid-like structure, GMpvs; grey matter perivascular space, WMpvs; white matter perivascular space.

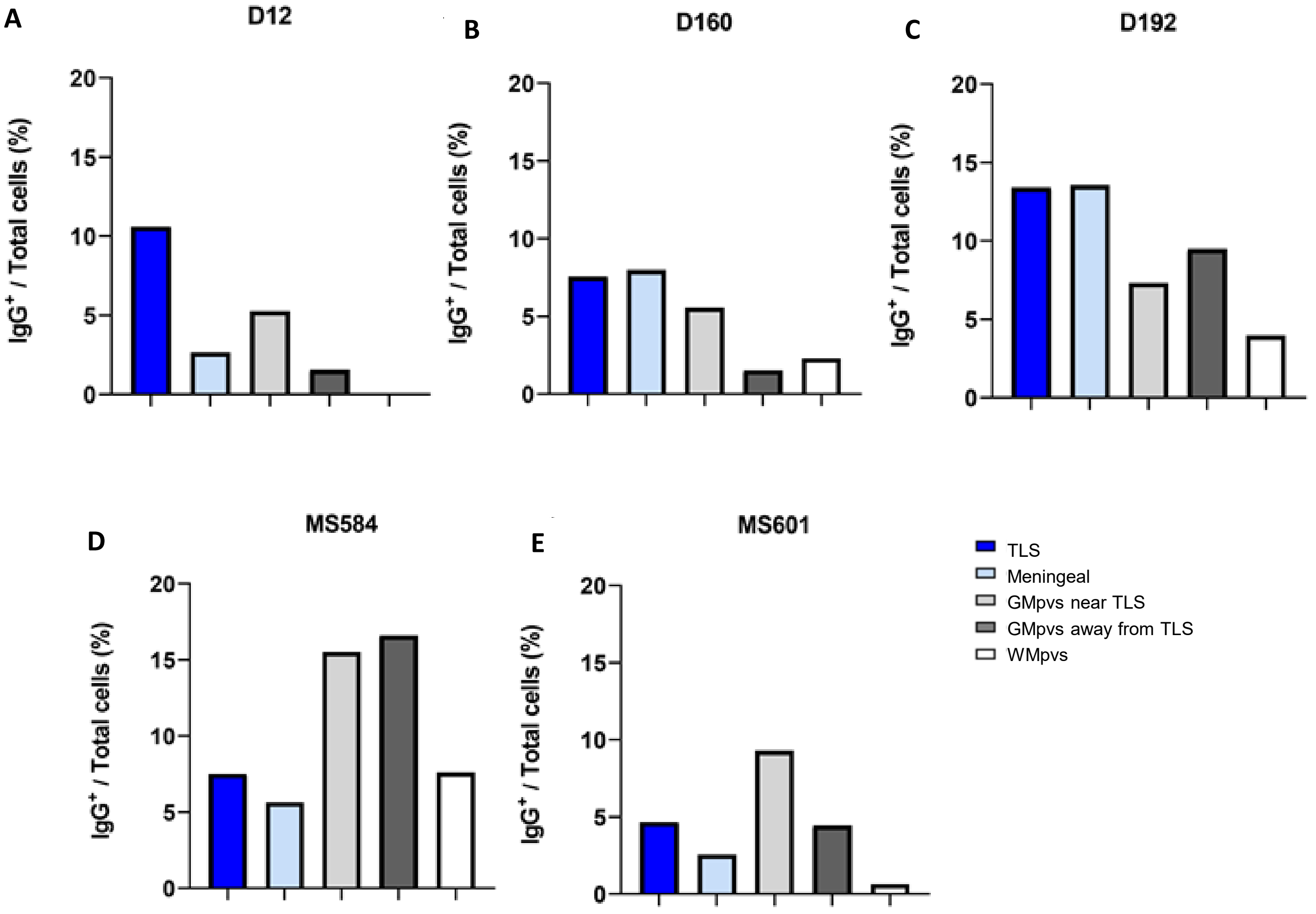


**Supplementary Figure 4: IgG$^+$ cell density in individual post-mortem cases.** Variation in IgG$^+$ cell density was observed across post-mortem cases. Higher IgG$^+$ cell densities were observed in TLS and other meningeal regions in case D160 and D192 (B, C). GM perivascular spaces in case MS584 and MS601 demonstrated higher IgG cell densities than other regions (D, E). Abbreviations: TLS; tertiary lymphoid-like structure, GMpvs; grey matter perivascular space, WMpvs; white matter perivascular space.

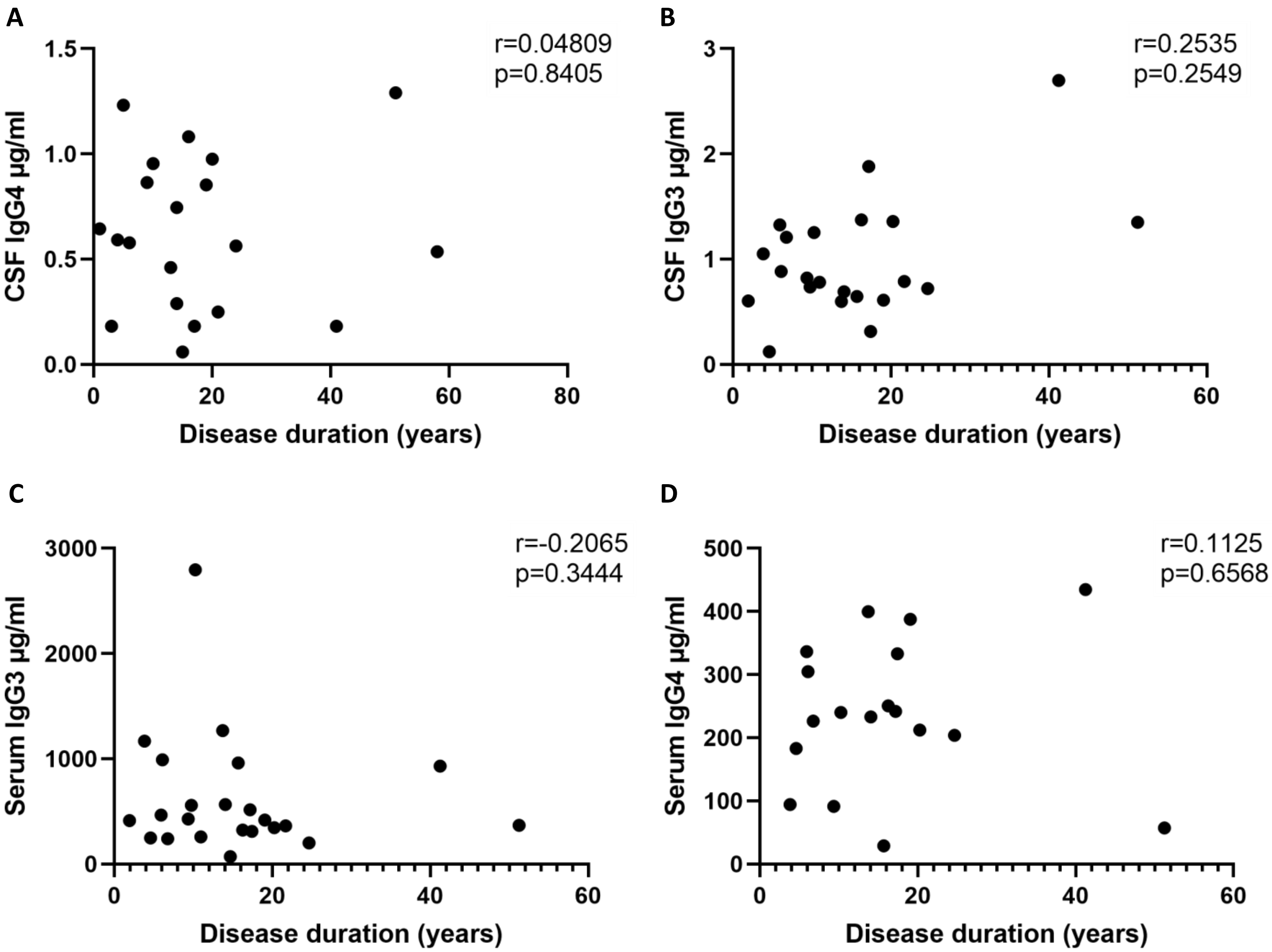


**Supplementary Figure 5: Serum and CSF IgG3 and IgG4 profiles.** Correlation plots showing relationship between serum and CSF IgG3 and IgG4 profiles with disease duration (A-D). Spearman correlation (serum and CSF IgG3 and disease duration). Pearson correlation (serum and CSF IgG4 and disease duration).

| Case ID | Diagnosis | Gender | Age at onset (years) | Age at death (years) | Disease duration (years) | Treatment with immune suppressant | Cause of death | PMI |
|---|---|---|---|---|---|---|---|---|
| D12 | Progressive | Male | 33 | 37 | 4 | None | Cardiac arrest | 14hrs |
| D25 | Progressive | Female | 26 | 30 | 4 | None | Not stated | 1 hr |
| D28 | PPMS | Male | 44 | 50 | 6 | None | Bronchopneumonia | <12hrs |
| D30 | SPMS | Female | 26 | 33 | 7 | None | Bronchopneumonia, metastatic thyroid cancer | <12hrs |
| D34 | PPMS | Male | 51 | 55 | 4 | None | Bronchopneumonia | <12hrs |
| D40 | PPMS | Female | 32 | 39 | 7 | None | Bronchopneumonia | 24hrs |
| D51 | MS | Female | 13 | 62 | 49 | None | Cancer | 24hrs |
| D78 | Progressive | Male | 52 | 56 | 4 | None | Bronchopneumonia | Not known |
| D86 | MS | Female | 49 | 52 | 3 | None | Road traffic accident | 25hrs |
| D112 | Benign MS | Female | Unknown | 81 | Unknown | None | Perforated duodenal ulcer, peritonitis | 24hrs |
| D131 | SPMS | Female | 28 | 71 | 43 | None | Pulmonary embolus | 4.3hrs |
| D134 | RRMS | Female | 37 | 41 | 3 | None | Bronchopneumonia, pulmonary embolus | Not known |
| D141 | SPMS | Female | 19 | 39 | 20 | None | Hypoxia | 24hrs |
| D142 | Benign MS | Male | 34 | 84 | 50 | None | Stroke | 7.75hrs |
| D144 | Progressive | Male | 47 | 54 | 7 | None | Bronchopneumonia | <12hrs |
| D145 | SPMS | Female | 26 | 66 | 40 | None | Bronchopneumonia | <12hrs |
| D147 | RRMS | Female | 30 | 33 | 3 | ACTH | Bronchopneumonia, pulmonary embolus | >24hrs |
| D153 | SPMS | Female | 26 | 50 | 24 | None | Pulmonary oedema | 48hrs |
| D158 | SPMS | Female | 34 | 62 | 28 | None | Bronchopneumonia | 24hrs |
| D159 | SPMS | Female | 32 | 76 | 44 | None | Pulmonary oedema, duodenal perforation | 1.75hrs |
| D160 | RRMS | Male | 47 | 51 | 2 | Hydrocortisone | Bronchopneumonia | 24hrs |
| D162 | SPMS | Male | 23 | 70 | 47 | None | Lung cancer | 1.25hrs |
| D163 | SPMS | Male | 53 | 70 | 17 | ACTH | Bronchopneumonia, myocardial infarction | 5.5hrs |
| D164 | Progressive | Female | Unknown | 62 | Unknown | None | Pulmonary infarction, pulmonary embolus | 1.50hrs |
| D165 | MS | Female | Unknown | 60 | Unknown | None | Bronchopneumonia | 4.75hrs |
| D166 | SPMS | Male | 25 | 46 | 21 | None | Pulmonary embolus | 24hrs |
| D168 | SPMS | Female | 52 | 82 | 30 | None | Bronchopneumonia | 24hrs |
| D170 | PPMS | Female | 50 | 66 | 16 | None | Bronchopneumonia | 6hrs |
| D173 | SPMS | Female | 20 | 31 | 10 | None | Infarction anterior pituitary, cerebral oedema | <24hrs |
| D174 | SPMS | Male | 43 | 58 | 15 | None | Myocardial infarction | 27hrs |
| D175 | SPMS | Female | 30 | 53 | 23 | None | Bronchopneumonia | 24hrs |
| D178 | PPMS | Male | 34 | 59 | 25 | None | Bronchopneumonia | 24hrs |
| D185 | SPMS | Female | 41 | 61 | 20 | None | Bronchopneumonia | <24hrs |
| D186 | Progressive | Female | 32 | 54 | 22 | None | Bronchopneumonia | <24hrs |
| D187 | SPMS | Female | 14 | 45 | 31 | None | Bronchopneumonia | 24hrs |
| D192 | RRMS | Female | 22 | 22 | 0.33 | ACTH | Hypoxic injury | 24hrs |
| D199 | PPMS | Female | 37 | 75 | 38 | None | Pyelonephritis | 24hrs |
| D202 | SPMS | Female | 25 | 35 | 10 | Steroids (>3 months from death) | Peritonitis | 24hrs |
| D206 | SPMS | Male | 49 | 65 | 16 | None | Pulmonary embolus, lung cancer | 24hrs |
| D207 | SPMS | Male | 23 | 34 | 11 | None | Bronchopneumonia | 1.3hrs |
| D211 | SPMS | Female | 40 | 55 | 15 | None | Bronchopneumonia | 9hrs |
| D213 | SPMS | Male | 24 | 33 | 9 | ACTH | Bronchopneumonia | 7.5hrs |
| D215 | RRMS | Male | 24 | 33 | 9 | ACTH | Septicaemia | 4.5hrs |
| D230 | Progressive | Female | 34 | 59 | 25 | None | Bronchopneumonia | <12hrs |
| D243 | SPMS | Female | 41 | 80 | 39 | None | Bronchopneumonia | 48hrs |
| D248 | RRMS | Female | 39 | 40 | 1 | Steroids, cyclophosphamide | Cerebral oedema | 24hrs |
| D253 | CIS | Male | 61 | 64 | 3 | None | Myocardial infarction | 24hrs |
| D261 | RRMS | Female | 22 | 22 | 0.5 | Methylprednisolone | Bronchopneumonia | 12hrs |
| D276 | SPMS | Female | 23 | 40 | 17 | None | Pneumonitis | Not known |
| D277 | SPMS | Female | 24 | 40 | 16 | None | Bronchopneumonia | <24hrs |
| D278 | SPMS | Male | 36 | 72 | 36 | None | Intracerebral haemorrhage | <24hrs |
| D279 | SPMS | Female | 28 | 64 | 36 | None | Peritonitis | 24hrs |
| D285 | SPMS | Female | 44 | 70 | 26 | None | Bronchopneumonia | <24hrs |

**Supplementary Table 1: Demographics of complete cohort examined for meningeal and perivascular inflammation.** Abbreviations: ACTH, adrenocorticotropic hormone. CIS, clinically isolated syndrome, MS, multiples sclerosis, PPMS, primary progressive MS, SPMS, secondary progressive MS, PMI, post-mortem interval.

| Measure | Mild Meningeal Score (0/+) | Moderate Meningeal Score (++) | Substantial Meningeal Score (+++) | p value |
|---|---|---|---|---|
| Cases (%) | 27 (54) | 16 (32) | 7 (14) | |
| Female (n) | 19 | 10 | 4 | |
| Male (n) | 8 | 6 | 3 | |
| Gender ratio (F/M) | 2.38 | 1.67 | 1.33 | |
| Age at onset (years) | 37 ± 9.75 | 30 ± 5.75 | 33 ± 10.5 | 0.4378 |
| Disease duration (years) | 16.5 ± 11.25 | 19.5 ± 10.75 | 4 ± 2 | 0.0186 |
| Age at death (years) | 60 ± 12.25 | 56 ± 12 | 37 ± 8.25 | 0.0511 |

**Supplementary Table 2: Characteristics of cases with mild, moderate and substantial meningeal inflammation scores.** Median ± half interquartile range is reported for age at onset, disease duration and age at death. Differences were tested for statistical significance by one-way ANOVA.

#### Supplementary Table 3: RNAscope HiPlex assay probes

| Probe | Target | Cat. No |
|---|---|---|
| RNAscope® Probe - Hs-IGHA1-T1 | IgA | 571211-T1 |
| RNAscope® HiPlex Probe - Hs-IgG-pan-T10 | IgG | 439501-T10 |
| RNAscope® HiPlex Probe - Hs-IGHM-T11 | IgM | 481091-T11 |
| RNAscope HiPlex12 Positive Control Probe v2-Rn | RTU mixture of 12 probes targeting housekeeping gene Polr2a, PPIB, UBC, HPRT1, Actb, Sdha, Tfrc, Ldha, Gapdh, Rpl5, Ywhaz, and Rpl28 with T1- T12 tails respectively in each of the 12 channels | 324434 |
| RNAscope HiPlex12 Negative Control Probe | RTU probe targeting a bacterial gene (dapB), with T1- T12 tails respectively in each of the 12 channels | 324301 |

#### Supplementary Table 4: Primary antibodies

| Antibody | Target | Host | Dilution | Clone | Supplier/Cat. No. |
|---|---|---|---|---|---|
| Anti-IgG | Human IgG | Rabbit | 1:500 | Monoclonal EPR4421 | Abcam Ab109489 |
| Anti-IgG3 | Heavy chain human IgG3 | Rabbit | 1:200 | Monoclonal RM119 | Abcam Ab193172 |
| Anti-IgG4 | Human IgG4 | Mouse | 1:100 | Monoclonal IHG4/1345 | Abcam Ab218428 |
| Anti-CD38 | Germinal centre B cells, plasma cells | Mouse | 1:100 | Monoclonal SPC32 | Leica CD38-290-L-CE |
| Anti-CD38 | Germinal centre B cells, plasma cells | Rabbit | 1:100 | Monoclonal SP149 | Merck SAB5500063 |
| Anti-CD138 | Plasma cells, keratinocytes, fibroblasts, endothelial cells, vascular smooth muscle cells | Goat | 1:100 | Polyclonal | Thermofisher PA547395 |

| Case ID | Cohort | Diagnosis | Gender | Age at LP (years) | Time from symptom onset to LP (years) | Disease duration at follow up (years) | EDSS: at time of LP | EDSS: at follow up |
|---|---|---|---|---|---|---|---|---|
| 44299 | RRMS at follow up | RRMS | M | 36 | 13.6 | 20.3 | 2 | <4.0 |
| 83643 | RRMS at follow up | RRMS | F | 36 | 5.0 | 9.8 | 0 | <4.0 |
| 11953 | RRMS at follow up | RRMS | F | 40 | 0.1 | 15.7 | <4.0 | <4.0 |
| 27835 | RRMS at follow up | RRMS | F | 23 | 6.1 | 14.1 | 1 | <4.0 |
| 28012 | RRMS at follow up | RRMS | F | 28 | 10.8 | 19.1 | <4.0 | <4.0 |
| 43475 | RRMS at follow up | RRMS | F | 23 | 3.1 | 10.3 | 2 | 1.5 |
| 13368 | RRMS at follow up | RRMS | M | 48 | 5.6 | 13.7 | 1 | <4.0 |
| 84390 | RRMS at follow up | RRMS | F | 45 | 1.4 | 6.1 | 1 | 2 |
| 12627 | RRMS at follow up | RRMS | F | 25 | 7.0 | 21.7 | 5.5 | <4.0 |
| 13229 | RRMS at follow up | RRMS | M | 24 | 1.1 | 14.7 | 1 | <4.0 |
| 97485 | RRMS at follow up | RRMS | F | 41 | 0.4 | 3.8 | 1.5 | 1 |
| 101879 | RRMS at follow up | RRMS | F | 48 | 2.0 | 4.6 | <4.0 | <4.0 |
| 121852 | RRMS at follow up | RRMS | F | 42 | 0.3 | 1.9 | 2 | <4.0 |
| 41964 | SPMS at follow up | SPMS | F | 51 | 2.1 | 9.4 | 4.5 | 6.5 |
| 46249 | SPMS at follow up | SPMS | F | 43 | 9.5 | 16.3 | <4.0 | 6 |
| 75244 | SPMS at follow up | SPMS | M | 70 | 45.6 | 51.3 | 3.5 | 6.5 |
| 12301 | SPMS at follow up | SPMS | M | 46 | 1.9 | 17.2 | 2.5 | 8.5 |
| 18166 | SPMS at follow up | SPMS | F | 65 | 1.1 | 11.0 | <4.0 | 5 |
| 11997 | SPMS at follow up | SPMS | F | 48 | 25.6 | 41.3 | 2.5 | 8 |
| 22789 | SPMS at follow up | SPMS | F | 47 | 8.3 | 17.4 | 0 | 6.5 |
| 21309 | SPMS at follow up | SPMS | M | 68 | 15.4 | 24.7 | 3.5 | 6.5 |
| 72829 | SPMS at follow up | SPMS | M | 36 | 0.1 | 5.9 | 3 | 5 |
| 109863 | SPMS at follow up | SPMS | F | 55 | 4.5 | 6.8 | 0 | 6 |
| 18771 | Control | IIH | M | 38 | n/a | n/a | n/a | n/a |
| 25236 | Control | Migraine | F | 21 | n/a | n/a | n/a | n/a |
| 26506 | Control | IIH | M | 43 | n/a | n/a | n/a | n/a |
| 27051 | Control | Migraine | M | 41 | n/a | n/a | n/a | n/a |
| 27976 | Control | Sensorineural hearing loss, functional disorder | M | 42 | n/a | n/a | n/a | n/a |
| 64974 | Control | Fibromyalgia | F | 57 | n/a | n/a | n/a | n/a |
| 66907 | Control | IIH | F | 23 | n/a | n/a | n/a | n/a |
| 67712 | Control | Migraine | F | 54 | n/a | n/a | n/a | n/a |
| 70485 | Control | IIH | F | 23 | n/a | n/a | n/a | n/a |

| | RRMS at follow up | | | | SPMS at follow up | | | |
|---|---|---|---|---|---|---|---|---|
| | median | min | max | SD | median | min | max | SD |
| **Age at LP (years)** | 36.0 | 23 | 48 | 9.2 | 49.5 | 36 | 70 | 10.8 |
| **Symptom onset to LP (years)** | 3.1 | 0.1 | 13.6 | 4.1 | 6.4 | 0.1 | 45.6 | 13.6 |
| **Disease duration at follow up (years)** | 13.7 | 1.9 | 21.7 | 6.2 | 16.7 | 5.9 | 51.3 | 14.3 |
| **EDSS: at time of LP** | 2 | 0 | 5.5 | | 3.25 | 0 | 4.5 | |
| **EDSS: at follow up** | <4.0 | 1 | <4.0 | | 6.5 | 5 | 8.5 | |

**Supplementary Table 5: Demographics of MS cohorts classified as RRMS or SPMS at follow up and non-MS control cohorts for paired serum and CSF samples.**

**Abbreviations:** IIH, idiopathic intracranial hypertension; LP, lumbar puncture; EDSS, expanded disability status scale; M=male; F=female
